# Insulin signaling gates long-term memory formation in *Drosophila* larvae

**DOI:** 10.1101/842997

**Authors:** Melanie Eschment, Hanna R. Franz, Nazlı Güllü, Luis G. Hölscher, Ko-Eun Huh, Annekathrin Widmann

## Abstract

The ability to learn new skills and to store them as memory entities is one of the most impressive features of higher evolved organisms. However, not all memories are created equal; some are short-lived forms, and some are longer lasting. Formation of the latter is energetically costly and by the reason of restricted availability of food or fluctuations in energy expanses, efficient metabolic homeostasis modulating different needs like survival, growth, reproduction, or investment in longer lasting memories is crucial. Whilst equipped with cellular and molecular pre-requisites for formation of a protein synthesis dependent long-term memory (LTM), its existence in the larval stage of *Drosophila* remains elusive. Considering it from the viewpoint that larval brain structures are completely rebuilt during metamorphosis, and that this process depends completely on accumulated energy stores formed during the larval stage, investing in LTM represents an unnecessary expenditure. However, as an alternative, *Drosophila* larvae are equipped with the capacity to form a protein synthesis independent so-called larval anaesthesia resistant memory (lARM), which is consolidated in terms of being insensitive to cold-shock treatments. Motivated by the fact that LTM formation causes an increase in energy uptake in *Drosophila* adults, we tested the idea of whether an energy surplus can induce the formation of LTM in the larval stage. Indeed, increasing the metabolic state by feeding *Drosophila* larvae the disaccharide sucrose directly before aversive olfactory conditioning led to the formation of a larval LTM (lLTM). Moreover, we show that the metabolic state acts as a binary switch between the formation of lARM and lLTM. Based on this finding, we determined that it is the insulin receptor (InR) expressed in the mushroom body Kenyon cells (MB KCs) that mediates this switch to favor the formation of lLTM under energy-rich circumstances and lARM under energy-poor circumstances.

## Introduction

Harboring the ability to deal with novelties and unpredictable complexities provides the key to successfully adapt to unforeseen events in an ever-changing environment. Therefore, one of the most outstanding capabilities if higher evolved organisms is the capacity to constantly learn new tasks, integrate new skills and preserve them as memory entities. However, establishing a memory is a highly complex and dynamic process. Apart from the involvement of multilayered neuronal circuitries and cellular machineries [1], the capacity to form memories comes with energetic costs since activation and maintenance of synaptic connections involved in integrating, storing and retrieving information are energy demanding [2,3]. These circumstances can either lead to trade-offs with other phenotypic traits or to learning and memory impairments, when available energy recourses are restricted [4,5]. For example, trade-offs between learning abilities in longevity and competitive abilities in *Drosophila* [6–8], reduced foraging skills in bumble bees [9], delayed juvenile development in mites [10], and decreased fecundity in guppies [11] and butterflies [12] have been described. Moreover, honeybees experience significant costs for learning and show a memory deficit being energetically stressed [13]. On the other hand, formation of LTM led to reduced resistance to food and water stress in *Drosophila* [14] and during food deprivation the formation of energetically costly LTM is disabled [15].

A general feature of memory formation across species is the parallel and chronologically ordered occurrence of distinct short-, intermediate-, and/or long-lasting memory phases [1]. In adult *Drosophila*, four temporally distinct memory phases have been characterized [16]. Thereby, LTM and ARM represent longer lasting memories that are resistant to anesthetic disruption but are mutually exclusive and distinguished by their dependence on *de novo* protein synthesis; LTM requires protein synthesis whereas ARM does not [17,18]. In adult *Drosophila* the formation of LTM, by protein synthesis dependency [17], causes an increase in energy uptake [19]. Under conditions of reduced food availability, the brain disables the formation of costly LTM and favors the formation of ARM [15]. One hypothesis proposes that “neuronal gating mechanisms” prevent adult *Drosophila* from forming energetically costly LTM under critical nutritional circumstances [19,20].

The larval stage of *Drosophila* has emerged as a favorable model system for studying learning and memory [21] because of the relative simplicity of the brain, for which the complete synaptic connectome is known [22,23]. Olfactory memory during the larval stage of *Drosophila* also consists of different memory phases [23]. For example, after classical aversive Pavlovian conditioning, during which larvae associate an odor with an aversive high salt stimulus [24,25], at least two co-existing memory phases have been distinguished: a labile larval short-term memory (lSTM) and lARM that are encoded by separate molecular pathways [24]. Although, *Drosophila* larvae possess cellular and molecular pre-requisites of potentially forming a protein synthesis dependent long-term memory (LTM) [23], the existence of a protein synthesis dependent LTM remains still elusive. Memorizing behavioral adjustments based on previous experience depends on the balancing of costs and benefits: only relevant information should be stored into energetically costly, protein synthesis dependent longer lasting memories, whereas less reliable information should be disregarded. Accordingly, the formation of LTM in larvae would represent an unnecessary expenditure, since larval brain structures are completely rebuilt during metamorphosis – meaning any plastic changes that occur due to learning might be lost in the re-wiring of the brain. Therefore, the aim of this study was to attempt to override this state-dependent limitation on LTM formation by feeding sugar prior to classical aversive conditioning. Indeed, we were able to show that by elevating the energetic state of larvae before conditioning, larvae are able to successfully form aversive lLTM. Conversely, we show that such a protocol inhibits the formation of lARM. We were additionally able to demonstrate that the process of lLTM formation depends on the activity of the *rutabaga* (*rut*) adenylate cyclase (AC), and that insulin receptors (IRs) expressed in the mushroom body Kenyon cells (MB KCs) gate the state-dependent switch between lARM and lLTM.

## Results

### Sucrose consumption specifically suppresses lARM

We first asked whether an increase in nutritional energy through carbohydrate uptake over a short period of time affects lARM. To tackle this question, we tested the memory performance of third instar, wild-type larvae trained using a previously described three-cycle aversive olfactory conditioning protocol [24], which was here additionally preceded by sucrose feeding for 60 min— to elevate the energetic state—and followed by an anesthetizing cold shock treatment (4°C) for 1 min [24]—to isolate lARM (Fig 1A and 1B). The memory tested 40 min after training onset (10 min after training offset) in larvae that consumed sucrose was indistinguishable from that of control larvae that consumed only tap water (Fig 1C, S3 Table). This memory was completely abolished after cold shock treatment (Fig 1C, S1 Table). Therefore, we concluded that lARM is not detectable after sucrose consumption anymore. It is unlikely that this memory phase is a residual lSTM, because it is well-established that lSTM is only detectable for up to 30 minutes after training onset using this aversive conditioning procedure [24]. Taking these findings into account, we hypothesize that sucrose consumption suppresses the expression of lARM.

**Fig 1.**
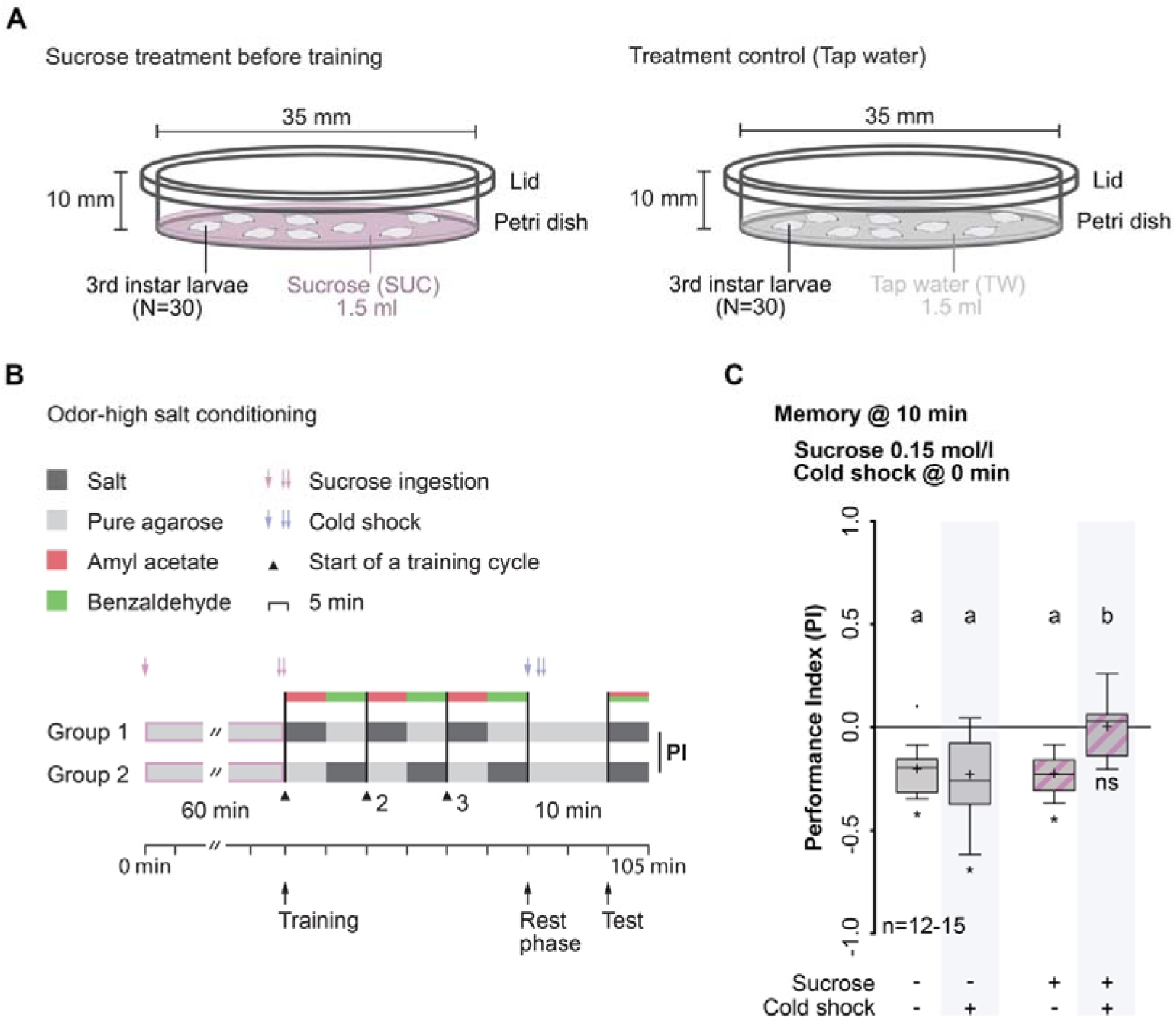
Sucrose consumption specifically suppresses lARM. (A) Schematic illustration of the manipulation of the metabolic state via sucrose feeding. Larvae were placed in a Petri dish filled with agarose containing either sucrose solution (SUC) (left) or tap water (TW) (right) for 60 min. (B) Schematic illustration of experimental principles and odor-high salt conditioning using a two-odor reciprocal training paradigm (reciprocally trained group not shown throughout). After feeding on 0.15 M SUC for 60 min (1 red arrow, start; 2 red arrows, end) two groups of 30 larvae were trained reciprocally with 3 training cycles without temporal gaps. Group 1 received the first odor AM paired with an aversive reinforcer (high salt concentration) while the second odor BA was presented alone (AM+/BA). Group 2 received the reverse contingency (AM/BA+). Subsequently, larvae received a cold shock treatment for 1 min (1 blue arrow, start; 2 blue arrows, end). Memory was tested 10 min later by calculating a Performance Index (PI). (C) After sucrose consumption, wild-type larvae showed a complete memory loss upon cold shock treatment. Larvae that consumed sucrose but did not receive a cold shock treatment showed lARM, comparable to larvae that did not consume sucrose independently of cold shock treatment. Memory performance above the level of chance was tested using Bonferroni-corrected one-sample t-tests (ns p≥0.0125; * p<0.0125; α=0.0125). Differences between the groups were determined using two-way ANOVA followed by Bonferroni *post-hoc* pairwise comparisons. Lowercase letters indicate differences between groups (p<0.05). For more statistical details see also Table S1 and S3. Data are shown as Tukey box plots; line, median; cross, mean; box, 75th-25th percentiles; whiskers, 1.5 interquartile range; small circles, outlier (n≥8). AM, n-amyl acetate; BA, benzaldehyde; lARM, larval anesthesia resistant memory; SUC, sucrose; TW, tap water.

Sugar consumption is regulated depending on the satiation state of the animal. In *Drosophila* larvae, hemolymph carbohydrate levels negatively correlate with sucrose consumption [26]. To ensure that this point of high sugar consumption was actually reached in our experiments, we examined the time at which sucrose consumption reached saturation by using a dye-feeding assay [27] (S1A Fig and S1B Fig). During the first 15 and 30 min, a steady increase in sucrose consumption was observed (S1A Fig and S1B Fig, S1 Table). By contrast, larvae feeding for 60 min showed sucrose ingestion behavior that was similar to that of larvae feeding on a dye-only solution (S1A Fig and S1B Fig, S1 Table), indicating that sucrose consumption had reached saturation within 60 min. Next, we confirmed that task-relevant sensory-motor abilities like naïve odor preference and salt avoidance were not altered after sucrose consumption (S1C Fig, S1 Table and S2 Table). Strikingly, the suppression of lARM after caloric intake was specific for sucrose and was an immediate effect, as neither the consumption of yeast for 60 min nor of high-caloric food for 1 day led to a suppression of lARM (S2A Fig and S2B Fig, S1 Table and S3 Table). This suggests the involvement of a fast-acting, specific sugar-detecting mechanism, rather than a general mechanism that monitors overall caloric food intake.

### Sucrose consumption gates a cAMP-dependent memory and inactivates *radish*-dependent lARM

The *radish* (*rsh*) gene [28] plays a pivotal role in the formation of lARM [24]. Using this mutant provides a tool to test whether the memory phase affected by sucrose consumption is equivalent to the molecularly defined lARM. In line with the key role of *rsh* in lARM formation [24], *rsh^1^* mutant larvae that fed on tap water for 60 min showed complete abolishment of an aversive olfactory memory tested directly after training, in contrast to wild-type animals (Fig 2A; S1 Table). However, the aversive olfactory memory of *rsh^1^* mutant larvae that consumed sucrose for 60 min prior to training revealed no significant defect in comparison with wild-type larvae that consumed either tap water or sucrose (Fig 2A; S3 Table). This finding suggests that the memory deficit in this ARM-specific memory mutant can be rescued by sucrose consumption. This further supports our hypothesis that lARM is replaced by an additional memory phase, if the energy state of the animal is sufficient. Next, we analyzed whether this rescue of memory in *rsh^1^* mutants is due to the direct action of sucrose in *rsh*-associated molecular pathways, or if there is an additional, *rsh*-independent mechanism at play. Again, we fed *rsh^1^* mutant larvae sucrose for 60 min, followed by conditioning and tested, if the formed aversive olfactory memory in these mutant larvae was sensitive to anesthesia induced by cold shock treatment (Fig 2B). No memory was detectable, indicating that the aversive olfactory memory formed in *rsh^1^* mutants after sucrose consumption was sensitive to cold shock treatment (Fig 2B, S1 Table).

**Fig 2.**
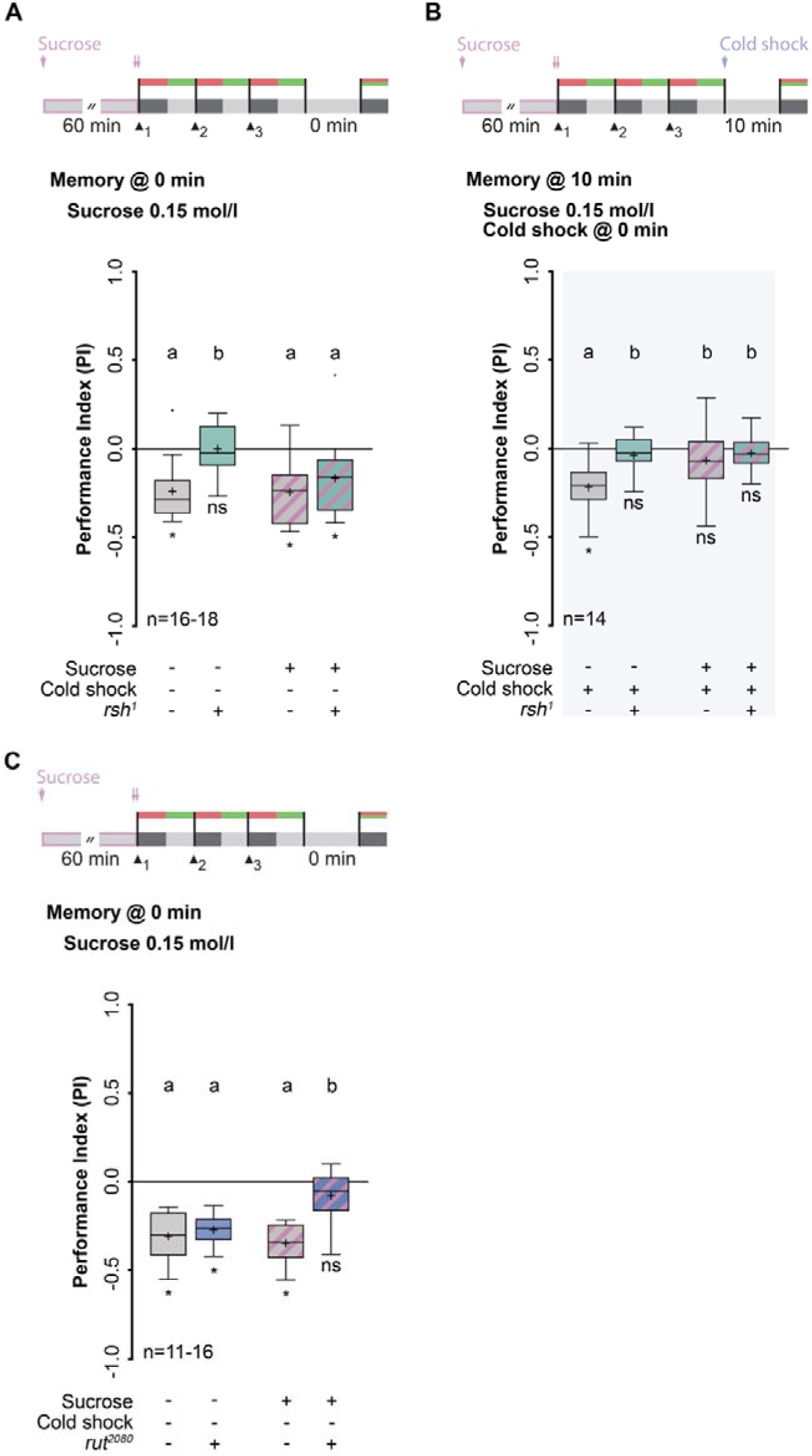
Sucrose consumption gates a cAMP-dependent memory and inactivates *radish*-dependent lARM. (A) Top: Training and treatment protocol. Memory was tested directly after training. Bottom: After sucrose consumption, memory formation is no longer impaired in *rsh^1^* mutants. (B) Top: Training and treatment protocol. Cold shock was applied to all groups. Memory was tested 10 min after training. Bottom: Memory in *rsh^1^* mutants after sucrose consumption is sensitive to cold shock treatment since they showed a complete memory loss. (C) Top: Training and treatment protocol. Memory was tested directly after training. Bottom: Sucrose consumption causes memory loss in *rut^2080^* mutants. Wild-type larvae fed either on tap water or sucrose and *rut^2080^* mutant larvae fed only on tap water showed memory formation indistinguishable from each other. Memory performance above the level of chance was tested using Bonferroni-corrected one-sample t-tests or Wilcoxon signed-rank test (ns p≥0.0125; * p<0.0125; α=0.0125). Differences between groups were determined using two-way ANOVA followed by Bonferroni *post-hoc* pairwise comparisons. Lowercase letters indicate differences between groups (p<0.05). For more statistical details see also Table S1 and S3. Data are shown as Tukey box plots; line, median; cross, mean; box, 75th-25th percentiles; whiskers, 1.5 interquartile range; small circles, outlier (n≥8). *rsh*, *radish*; *rut*, *rutabaga*.

Apparently, sugar consumption induces a memory phase that differs from lARM at the molecular level. Interestingly, previously reported genetic dissections of larval memory revealed that aversive lSTM and lARM utilize different molecular pathways [24,29,30], in which lSTM depends on proper cAMP-induced signaling. Therefore, we tested whether the formation of the cold shock-sensitive memory after sucrose consumption depends on cAMP signaling. We fed the classical learning mutant *rutabaga^2080^* (*rut^2080^*), which exhibits the inability to appropriately increase intracellular cAMP level [31], sucrose for 60 min followed by conditioning (Fig 2C). Directly after training, *rut^2080^* larvae fed on tap water showed intact aversive olfactory memory (Fig 2C, S1 Table and S3 Table), in line with the finding that *rsh*-dependent lARM, but not cAMP-dependent lSTM, is prevalent at this time point [4]. However, aversive olfactory memory after sucrose consumption in *rut^2080^* mutants was completely abolished (Fig 2C, S1 Table). These findings indicate that the newly formed aversive olfactory memory, induced through sucrose consumption, replaces *rsh*-dependent lARM with a *rut*-dependent memory. Therefore, sugar consumption triggers a switch between molecular pathways determining memory phases.

### Activity of the insulin receptor is necessary for suppression of lARM after sucrose consumption

In *Drosophila,* the insulin-like growth factor signaling (IIS) pathway is not only essential for maintaining energy storage and glucose metabolism, but also for regulating lifespan and aging, reproduction, nutrient sensing, and cellular growth [32]. In contrast to mammals, which have a large family of IIS-receptors, *Drosophila* has only one insulin receptor (DInR), but eight insulin-like peptides [33,34]. It has been shown that the DInR is necessary for the formation of aversive olfactory LTM in adult *Drosophila* and for the formation of intermediate-term memory in aged flies [35,36]. DInR is strongly expressed in the larval central nervous system (CNS), as well as that of adults [37]. Furthermore, mapping the developmental expression atlas of genes in MB neurons revealed an expression of the DInR in the MB of *Drosophila* third instar larvae [38]. This is important because the synapses that change in the course of associative olfactory learning and thereby mediate memory formation could be localized to the MB KCs, both in adult and larval *Drosophila* [23,39]. Thus, we tested whether the suppression of lARM after sucrose consumption depends on proper insulin signaling in the MB of larval *Drosophila*. We expressed a dominant negative variant of the DInR (UAS-*InR^DN^*) in all KCs using the driver line OK107. We fed OK107/UAS-*InR^DN^* and both control groups (OK107/+ and UAS-*InR^DN^*/+) sucrose for 60 min followed by conditioning and cold shock treatment (Fig 3A). Both control groups receiving cold shock treatment showed a complete abolishment of aversive olfactory memory after sucrose consumption (Fig 3A, S1 Table). By contrast, larvae expressing *InR^DN^* in KCs (OK107/UAS-*InR^DN^*) showed an intact aversive olfactory memory comparable to that of the three genetic groups (OK107/+, UAS-*InR^DN^*/+ and OK107/UAS-*InR^DN^*) that did not receive any cold shock after conditioning (Fig 3A, S1 Table and S3 Table). All task-relevant sensory-motor abilities were unaltered after sucrose consumption (S3A Fig – S3D Fig, S1 Table and S3 Table); however, larvae expressing *InR^DN^* in KCs (OK107/UAS-*InR^DN^*) showed a slight reduction in sucrose consumption (S3E Fig – S3G Fig, S1 Table). This is in line with the observation that inhibition of insulin signaling in the neurons of the MB reduces food intake [40]. Overall, we conclude that intact insulin signaling is necessary for the suppression of lARM and for the observed switch in memory phases after sucrose consumption. But which memory phase, exactly, is induced through sucrose consumption?

**Fig 3.**
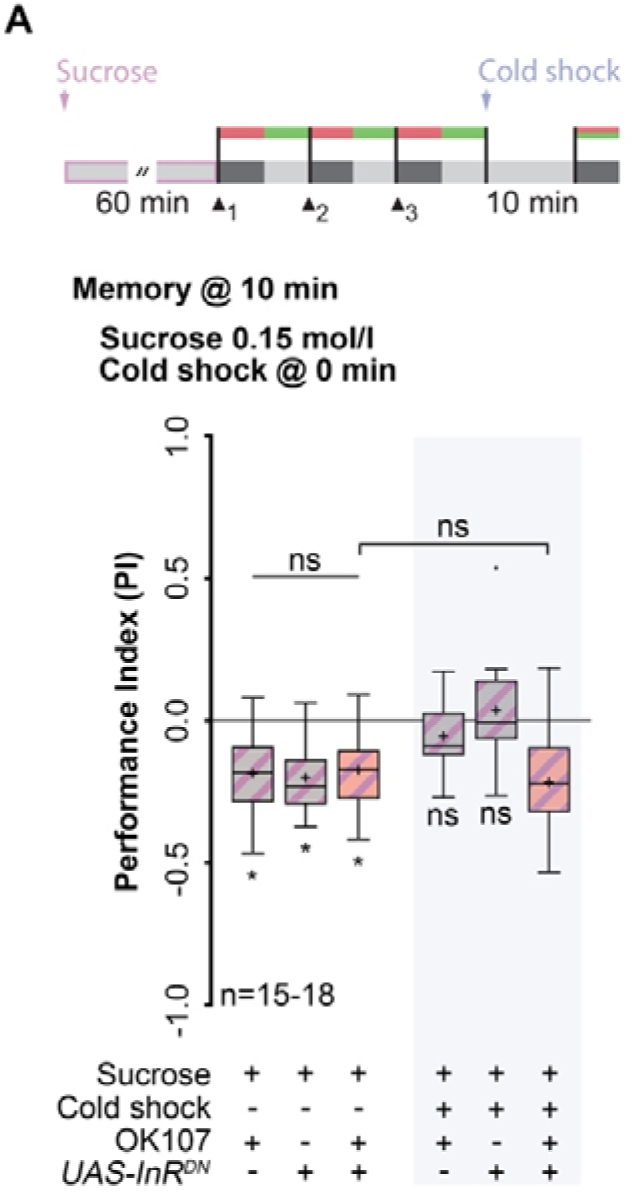
Activity of the insulin receptor is necessary for suppression of lARM after sucrose consumption. (A) Top: Training and treatment protocols. All groups consumed sucrose for 60 min and identification of lARM was carried out by applying a cold shock directly after training. Memory was tested 10 min after training. Bottom: Expression of the dominant negative form of the insulin receptor (UAS-*InR^DN^*) in KCs using the driver line OK107 prevents the suppression of lARM formation triggered by sucrose consumption. Memory performance above the level of chance was tested using Bonferroni-corrected one-sample t-tests (ns p≥0.008; * p<0.008; α=0.008). Differences between the groups were determined using two-way ANOVA followed by Bonferroni *post-hoc* pairwise comparisons. Statistically non-significant differences between groups (p≥0.05) are indicated as ns. For more statistical details see also Table S1 and S3. Data are shown as Tukey box plots; line, median; cross, mean; box, 75th-25th percentiles; whiskers, 1.5 interquartile range; small circles, outlier (n≥8). DN, dominant negative; InR, insulin receptor; KC, Kenyon cell; lARM, larval anesthesia resistant memory; UAS, upstream activation sequence.

### Rapid consolidation of lLTM after the consumption of sucrose

We have shown, that after sucrose consumption lARM is suppressed and a second, cAMP dependent memory component is formed (Fig 1B and 2C). But which memory phase, exactly, is induced through sucrose consumption? It has been shown in *Drosophila* adults that STM but also LTM rely on proper cAMP signaling [41]. So far, a protein synthesis-dependent LTM has not yet been shown in larvae, although evidence of a longer form of memory dependent on the transcription factor CREB strongly points towards its existence [24].Therefore, we questioned whether feeding on sucrose induces the formation of lLTM thereby switching lARM to lLTM. First, we determined whether the memory was stable over a longer period of time. We fed wild-type larvae sucrose for 60 min prior to conditioning and tested the memory 60 min after training (S4A Fig). The observed olfactory memory was found to be more stable than lSTM, based on the fact that it was still detectable after 60 min and was as robust as lARM formed without sucrose feeding (S4A Fig, S1 Table and S2 Table). Therefore, the newly formed memory was long-lasting on a larval time scale. Next, we tested whether the memory formed after sucrose consumption was dependent on *de-novo* protein synthesis by feeding larvae with the translation-inhibitor cycloheximide (CXM) for 16 hours before the sucrose feeding [17,24] (Fig 4A). Wild-type larvae treated with CXM and fed sucrose showed a statistically significant decrease in olfactory aversive memory tested at 30 min and 60 min after conditioning when compared to control groups, with the effect being stronger at 60 min (Fig 4B, S1 Table and S3 Table). However, the memory was not completely abolished (Fig 4B, S1 Table and S3 Table). These findings indicate that sucrose consumption leads to the suppression of lARM and, instead, promotes a rapid consolidation of larval LTM (lLTM).

**Fig 4.**
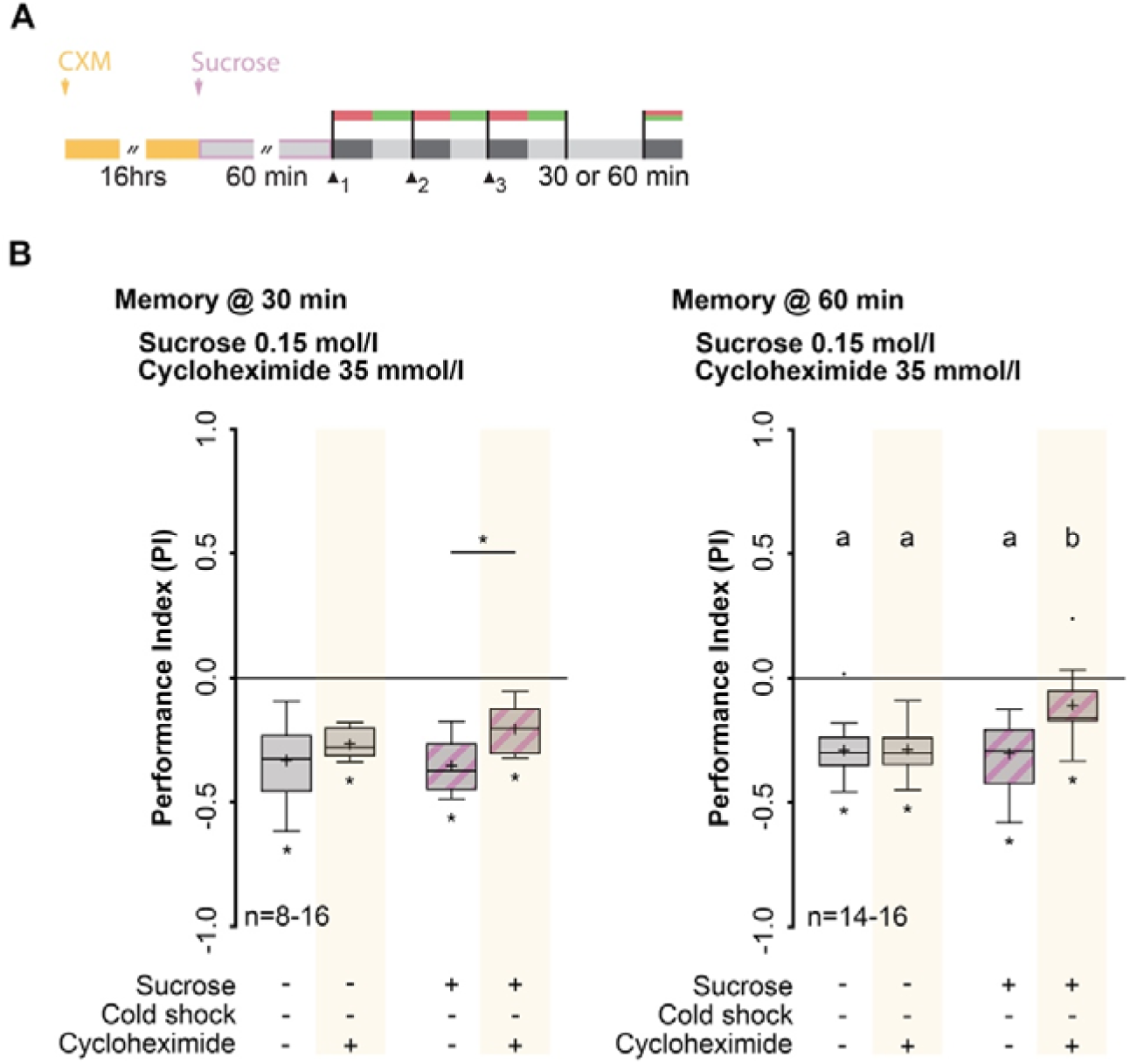
Rapid consolidation of lLTM after the consumption of sucrose. (A) Training and treatment protocols. Before feeding on sucrose, larvae were fed for 16 hours on CXM. Memory was tested 30 and 60 min after training. (B) Sucrose consumption gates rapid formation of a protein synthesis-dependent lLTM. Memory tested 30 min after training was only statistically different between larvae that consumed sucrose with or without CXM treatment. After CXM treatment, larvae that consumed sucrose showed only a slight memory 60 min after training, which was statistically significant different to all other groups of larvae. However, it was not completely abolished. Memory performance above the level of chance 30 was tested using Bonferroni-corrected one-sample t-tests (ns p≥0.0125; * p<0.0125; α=0.0125). Differences between groups were determined using two-way ANOVA followed by Bonferroni *post-hoc* pairwise comparisons. Lowercase letters indicate differences between groups (p<0.05). For more statistical details see also Table S1 and S3. Data are shown as Tukey box plots; line, median; cross, mean; box, 75th-25th percentiles; whiskers, 1.5 interquartile range; small circles, outlier (n≥8). CXM, cycloheximide; lLTM, larval long-term memory.

Typically, aversive olfactory LTM is induced by multiple training trials that are separated by temporal spaces [17]. In larvae, five spaced cycles of training leads to CREB-dependent lLTM [24]. We tested whether the sugar-induced formation of lLTM shown here matched the time course of spaced training-induced lLTM. After three spaced training trials, no lLTM was observed (S4B Fig, S1 Table and S2 Table). This result was in contrast to sugar-promoted lLTM formation, shown here to be inducible even after only three training trials. Therefore, we postulate that sugar gates lLTM formation more rapidly and efficiently than increasing the number of spaced training cycles. However, it has been shown that blocking protein synthesis using CXM has a deleterious effect over a longer period of time; specifically, larvae do not properly pupate or enclose [24]. Therefore, we tested whether sucrose consumption after CXM treatment was impaired by feeding larvae CXM for 16 hours. Larvae consumed a detectable amount of liquid dye (0.091±0.025 μl/larva/h, one-sample t-test, p=0.002) (S2 Data) and the consumption of sucrose was not altered after CXM treatment (S4C Fig, S1 Table). Therefore, the effect of CXM on memory formation cannot be attributed to impaired sucrose consumption.

## Discussion

Establishing a memory requires the timely controlled action of different neuronal circuits, neurotransmitters, neuromodulators and molecules. It is known that after classical aversive olfactory conditioning, *Drosophila* adults form two mutually exclusive longer lasting memory types -LTM and ARM -which can be distinguished based on their dependency on *de-novo* protein synthesis [17,18,42]. The occurrence of such genetically and functionally distinct memory phases is conserved in the animal kingdom, shown in honeybees, in *Aplysia*, and also in vertebrates [43–46].

The hypothesis that protein synthesis-dependent LTM formation is energetically costly and, therefore, restricted to favorable nutritional conditions, is based on a study of adult *Drosophila* [15]. Furthermore, after a spaced training protocol known to induce LTM formation [17,47], flies increased their sugar consumption [19]. Therefore, it seems that the formation of LTM is closely related to energy metabolism, such that the cost of this process must be compensated with increased sugar consumption. Larval *Drosophila* undergoes metamorphosis and the accumulated energy storage during this stage contributes to somatic maintenance and reproduction in adults [48]. Therefore, these larvae present a model system in which the energetic cost of LTM formation far exceeds the potential benefit, especially considering that this memory faces potential degradation during metamorphosis.

Seen in this light, and along with fact that larvae possess all the necessary cellular machinery, we hypothesized that short-term feeding on sucrose directly before training could result in a surplus of energy such that LTM formation is induced instead of ARM. Indeed, we show here that feeding larvae sugar before conditioning is also sufficient to trigger this switch, even with a less intensive training protocol. This implies that LTM formation is based on two gating mechanisms: one responding to the training intensity (e.g., temporal spacing of multiple trials) and one to the metabolic state. Regarding the first, two slow oscillating dopaminergic neurons have been proposed to act as a gating mechanism for LTM formation at the cost of inhibiting protein synthesis-independent ARM [20]. Regarding the latter, we propose a mechanism in the brain of larval *Drosophila* that directly senses the metabolic state at the time of training and is furthermore independent of the training regime (**Fig 5**). Without feeding on sucrose or by knocking down the InR in the MB KCs, two co-existing memory phases are visible after aversive olfactory conditioning (lSTM and lARM, Fig 5A). However, by elevating the energetic state by feeding sucrose and through an insulin-signaling-dependent gating mechanism, the *rsh*-dependent lARM is suppressed and a cAMP-dependent lLTM is visible (Fig 5B). This supports the finding that *Drosophila* larvae can form a CREB-dependent memory [24]. Therefore, we have determined that the conserved principal of cAMP-dependent, protein synthesis-dependent LTM formation holds true also for *Drosophila* larvae, although they undergo metamorphosis and most likely all formed lLTM is erased after the re-structuring of brain connectivity in the course of pupation.

**Fig 5.**
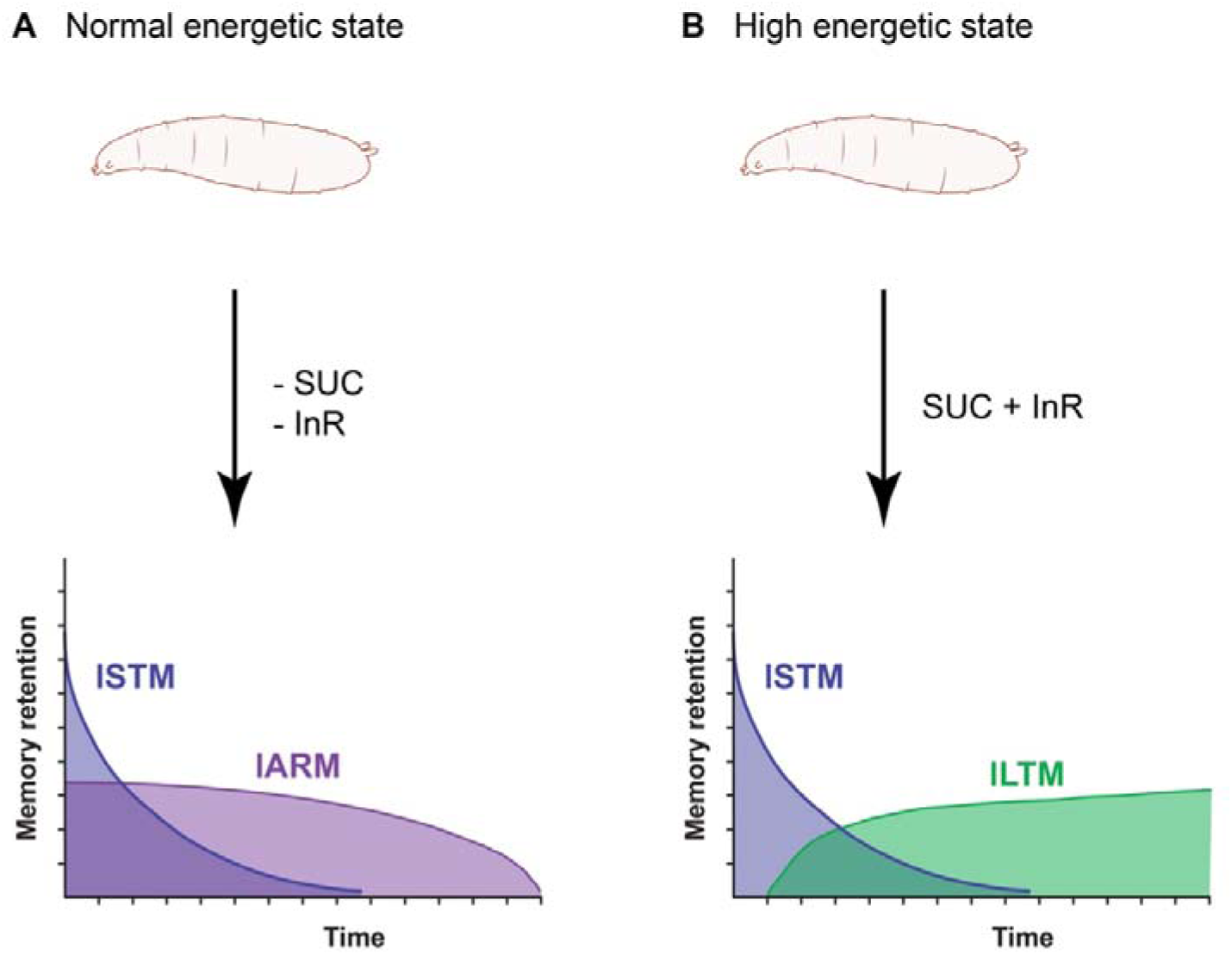
Working hypothesis on the state-dependent switch between lLTM and lARM after elevating the energetic state of *Drosophila* larvae. (A) In the absence of sucrose or by downregulation of the InR in the MB KCs *Drosophila* larvae form lSTM and lARM. (B) By feeding sucrose prior to conditioning lLTM formation instead of lARM is triggered, and this is dependent on the activity of the InR in the MB KCs. lARM, larval anesthesia resistant memory; lLTM, larval long-term memory; lSTM, larval short-term memory; InR, insulin receptor; SUC, sucrose.

Remarkably, we have also demonstrated a novel gating mechanism underlying the formation of LTM. Previous work has shown that LTM in adult *Drosophila* leads to a subsequent increase in energy metabolism. We take this a step further by demonstrating that increasing the energetic state of larvae before the training begins is sufficient to trigger the formation of LTM even after a less intense training protocol. This means that, although LTM is highly costly (and in the case of larvae theoretically redundant), its formation can be forced under the right circumstances due to the presence of a mechanism for the detection of energetic surplus that negates this high cost. This is also in agreement with recent studies showing that glycolytic enzymes are required in the MB of adult *Drosophila* for the formation of aversive olfactory memory [49]. This means that the brain of larval *Drosophila*—and potentially brains of other animals as well [50]—is not only a calculation device to decide if incoming sensory information is of importance, for example in the case of repetitions of the same stimulus, but can also sense and balance existing resources and decide if forming an expensive memory is an affordable or life-threatening luxury, especially for larvae whose main behavioral activity is taking in food.

Sugar by itself is assumed to be a primary source of energy, with circulating sugar levels reflecting the energetic state of an animal. Controlling the metabolic homeostasis is regulated via the *Drosophila* orthologs of glucagon (adipokinetic hormone, AKH) and insulin (*Drosophila* insulin-like peptide, DILP) [51]. Both have been proven to be involved in feeding and foraging behaviors and are controlled contrastingly through glucose [52–54]. Additionally, it has been shown that the InR is acutely required for LTM formation in *Drosophila* adults [36]. Strikingly, we show here that both increase in energetic state and the InR are necessary to mediate the formation of lLTM in *Drosophila* larvae (Fig 5B). We concluded that the InR in the MB KCs of *Drosophila* larvae can directly sense the elevated energetic state provoked by feeding sucrose directly before training and as a result mediate the state-dependent switch between lARM and lLTM. Beyond that, the involvement of insulin signaling in memory formation has striking parallels in mammals as well. For example, downregulation of an insulin receptor in the hippocampus of mice leads to spatial learning deficits [55]. Moreover, injections of insulin reversed memory deficits caused by Alzheimer’s disease, and in stroke patients an intranasal insulin treatment has been shown to improve hippocampal-dependent declarative memory in healthy humans [56]. Given the fact that the molecular underpinnings of both memory formation and insulin signaling are highly conserved across the animal kingdom [57,58], this correspondence among taxa is not surprising. Rather, it corroborates the general validity of model organisms like *Drosophila*. Thus, our finding that insulin signaling gates the formation of LTM and inhibits an alternative memory component could be of importance for the study in higher organisms, including humans.

## Acknowledgment

We are grateful to André Fiala and Andreas S. Thum for support. We thank Lyubov Pankevych, Margarete Ehrenfried and Jutta Böker for fly care and maintenance. Furthermore, we thank Claudius Neumann and Clare Hancock for proofreading the manuscript. We would like to thank Bertram Gerber, Thomas Preat, André Fiala and Andreas S. Thum for helpful comments on the manuscript and discussions. This work was supported by the University of Göttingen and the Zukunftskolleg of the University of Konstanz.

## Authors Contributions

Conceptualization: A.W. Data curation: A.W. Formal analysis: N.G., M.E., K.-E.H., H.R.F., L.H., A.W. Experimental work: N.G., M.E., K.-E.H., H.R.F., L.H., A.W. Project administration and supervision: A.W. Writing: A.W.

## Material and methods

### Fly stocks

Fly strains were reared on standard *Drosophila* medium at 25°C with 70% humidity in a 12-hour light-dark cycle. Crosses were raised at 18°C or 25°C with 70% relative humidity in a 12-hour light-dark cycle on standard *Drosophil*a medium. Flies were transferred to new vials and allowed to lay eggs for 2 days. For all experiments, 6-day-old foraging (feeding) third instar larvae were used. The wild-type strain was *Canton-S* (denoted here as wild-type). We used the learning mutants *rut^2080^* (obtained from the Bloomington Drosophila Stock Center, BDSC No.: 9405) and *rsh^1^* (kindly provided by T. Preat) [28,31]. All lines were outcrossed over several generations with wild-type *Canton-S* that was used as a genetic control. To express Gal4 in all larval Kenyon cells (KCs) we used the driver line OK107 [59,60] (obtained from the Bloomington Drosophila Stock Center, BDSC no.: 106098). The effector line UAS-*dInR^A1409K^* (denoted here as UAS-*dInR^DN^*) (obtained from the Bloomington Drosophila Stock Center, BDSC No.: 8253) was used to reduce insulin signaling within the KCs. The UAS-*dInR^DN^* transgene carries an amino acid replacement in the kinase domain (K1409A) of the *Drosophila* insulin receptor (dInR), which results in its dominant negative activity [61].

### Aversive olfactory learning and memory

Aversive olfactory learning and memory was performed at 23°C under standard laboratory conditions. Standard aversive olfactory conditioning experiments were performed using an odor-high salt conditioning paradigm, as previously described [24]. Experiments were conducted on assay plates (92-mm diameter, Sarstedt, Nümbrecht, cat. no.: 82.1472) filled with a thin layer of 2.5% agarose containing either pure agarose (Sigma Aldrich, cat. no.: A5093, CAS no.: 9012-36-6) or agarose plus 1.5 M sodium chloride (Sigma Aldrich, cat. no.: S7653, CAS no.: 7647-14-5) [24,25]. As olfactory stimuli, we used 10 μl amyl acetate (AM, Sigma Aldrich cat. no.: 109584; CAS No.: 628-63-7; diluted 1:250 in paraffin oil, Sigma Aldrich cat. no.: 18512, CAS no.: 8012-95-1) and benzaldehyde (BA, undiluted; Sigma Aldrich cat. no.: 418099, CAS no.: 100-52-7). Odorants were loaded into custom-made Teflon containers (4.5-mm diameter) with perforated lids [62]. Learning ability was tested by exposing a first group of 30 larvae to AM while they crawled on agarose medium that additionally contained sodium chloride as a negative reinforcer. After 5 min, the larvae were transferred to a fresh Petri dish in which they were allowed to crawl on a pure agarose medium for 5 min while being exposed to BA (AM+/BA). A second group of larvae received the reciprocal training (AM/BA+). Three training cycles were conducted. To test the memory after training, larvae were transferred onto another agarose plate and kept there for the indicated time before the memory was tested. To increase the humidity, tap water was added. Memory was tested by transferring larvae onto fresh agarose plates containing 1.5 M sodium chloride, on which AM and BA were presented on opposite sides. After 5 min, individuals located on the AM side (#AM), BA side (#BA), or in a 1-cm neutral zone were counted. We determined a preference index for each training group by subtracting the number of larvae on the BA side from the number of larvae on the AM side, and dividing by the total number of counted individuals (#TOTAL), as follows:

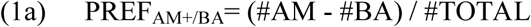

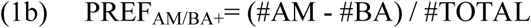

To specifically measure the effect of associative learning that is of the odor-reinforcement contingency, we then calculated the associative Performance Index (PI) as the difference in preference between the reciprocally trained larvae, as follows:

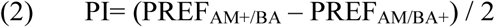

Negative PIs represented aversive associative learning, whereas positive PIs indicated appetitive associative learning. Division by 2 ensured that the scores were bounded between −1 and 1).

### Manipulation of the nutritional state

The nutritional state of larvae was manipulated by feeding 0.15 M sucrose (Sigma Aldrich, cat. no.: 84097, CAS no.: 57-50-1) for 60 min. A group of 30 larvae were either fed with 0.15 M sucrose mixed with tap water (+SUC) or with tap water (-SUC, control group). Larvae were placed in a Petri dish (35-mm diameter, Sarstedt, Nümbrecht, cat. no.: 82.1135.500) containing 2.5% agarose, and 1.5 ml of sucrose solution (+SUC) or tap water (-SUC) was added. This volume ensured that larvae did not crawl out of the sucrose solution and additionally prevented them from drowning. The larvae were allowed to feed for 60 min (if not stated otherwise) at 23°C. Zeitgeber time and humidity were kept constant for these experiments. The larvae were washed gently with tap water after being fed and transferred to an empty Petri dish containing 2.5% agarose.

### Quantification of sucrose consumption

To quantify sucrose consumption we used a modified feeding assay, as previously described [27]. A group of 30 larvae were placed in a Petri dish (35-mm diameter, Sarstedt, Nümbrecht, cat. no.: 82.1135.500) containing 2.5% agarose and either 1.5 ml 0.15 M sucrose + 2% indigo carmine (w/vol) (Sigma Aldrich, cat. no.: 57000, CAS no.: 860-22-0) mixed in tap water (+SUC +IC, experimental group), 2% (w/ml) indigo carmine mixed in tap water (+IC, dye-only control) or tap water (-IC, blank control). Again, 1.5 ml of the specific solution ensured that larvae did not crawl out of the solution and additionally prevented them from drowning. The larvae were allowed to feed for 1 hour at 23°C. Zeitgeber time and humidity were kept constant for these experiments. After 60 min, larvae were rinsed with tap water, transferred into 2-ml Eppendorf cups containing 500 μl of 1 M L-ascorbic acid (Sigma Aldrich, cat. no.: A7506, CAS no.: 50-81-7) and bead-based homogenized for 2 min using a Qiagen TissueLyser LT at a frequency of 50/s. After centrifugation at 14,800 rpm for 5 min at 23°C, the supernatant (400 μl) was transferred to Micro Bio-Spin Columns (Bio-Rad) and centrifuged again at 14,800 rpm for 5 min at 23°C for filtration. Subsequently, 200 μl of the supernatant was transferred into a new Eppendorf cup (1.5 ml) and centrifuged for a third time at 14,000 rpm for 2 min at 23°C. Two quantify sucrose consumption, 100 μl supernatant was transferred to a 96-well plate (Greiner Bio-One, cat. no.: 655061) and absorbance was measured at 610 nm [27] using a BioTek^TM^ Epoch Spectrophotometer. The corrected absorbance ABS (CORR) of each measurement was calculated by subtracting the mean absorbance of 1 M ascorbic acid (ABS_AA_) from the relative absorbance of either the blank control (ABS_-IC_), dye-only control (ABS_+IC_), or experimental group (ABS_+SUC +IC_). The relative consumption of sucrose (R.C.) was deduced by calculating the difference between the corrected mean absorbance of the blank control (ABS_-IC_(CORR)), the corrected mean absorbance of the dye-only control (ABS_+IC_(CORR)) and the relative absorbance of the experimental group (ABS_-IC +SUC_(CORR)):

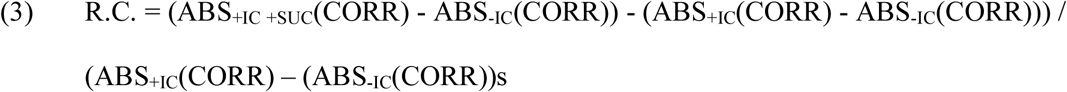

The blank control and the dye-only control were measured at every experiment and for every genotype on the same day. An R.C. value of 0 indicated that the larvae in the experimental group ate as much as the dye-only control larvae, a R.C.≤0 indicated that the larvae in the experimental group ate less than dye-only control larvae, and an R.C.≥0 indicated that larvae in the experimental group ate more than larvae in the dye-only control. To verify that the amount of ingested dye is represented in a linearly proportional manner, absorbance at 610 nm was measured for 100 μl of ascorbic acid and 2% (w/ml) indigo carmine in a two-fold serial dilution (data not shown).

### Cold shock treatment

To distinguish between cold shock sensitive and cold shock resistant memory phases, odor-high salt conditioning was followed by a cold shock treatment, as previously described [24]. Briefly, larvae were incubated in ice-cold tap water (4°C) for 1 min. Larvae were allowed to recover for at least 10 min by transferring them onto fresh agarose plates. They started moving within 2 min and were kept on the agarose plates at 23°C until testing.

### Cycloheximide treatment

To test if aversive olfactory memory induced by feeding sucrose prior to training is dependent on *de novo* protein synthesis, larvae were fed cycloheximide (CXM) as previously described [24]. Briefly, larvae were fed either with 35 mM cycloheximide (+CXM; Sigma Aldrich cat. no.: C7698; CAS no.: 66-81-9) or tap water (-CXM, control group) for 16 hours before the experiment. Therefore, 300 µl of CXM solution or tap water was added to the food vials. Before the experiment the larvae were gently washed with tap water and transferred to an empty Petri dish before being fed sucrose and undergoing subsequent odor-high salt conditioning and testing of the aversive olfactory memory at different time points.

### Odor preference and high salt avoidance experiments

To analyze larval olfactory perception, 30 larvae were placed along the midline of a Petri dish containing 2.5% pure agarose, with either a 10 μl amyl acetate-(AM) or a benzaldehyde-containing (BA) odor container on one side and an empty container (EC) on the other side. After 5 min, larvae located on the odor side (#ODOR), the side with the empty container (#EC), or in a 1-cm neutral zone were counted. By subtracting the number of larvae on the odor side from the number of larvae on the EC side, and dividing by the total number of counted individuals (#TOTAL), we determined a preference index for either AM or BA for each training group, as follows:

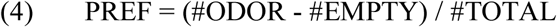

To investigate high salt avoidance, 30 larvae were placed along the midline of a Petri dish containing pure agarose on one side and agarose plus 1.5 M sodium chloride on the other. After 5 min larvae located on the salt side (#SALT), the agarose side (#AGAROSE), or in a 1-cm neutral zone were counted. By subtracting the number of larvae on the odor side from the number of larvae on the EC side, and dividing by the total number of counted individuals (#TOTAL), we determined a preference index for high salt avoidance for each training group, as follows:

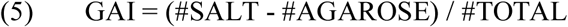

### Quantification and statistical analysis

All statistical analyses and visualizations were conducted with *GraphPad Prism* 8.0.2. Significance level of all statistical test was set to α=0.05. To compare single groups against the level of chance, we used Bonferroni-corrected two-tailed one-sample t-tests for normally distributed data (Shapiro-Wilk test), otherwise Bonferroni-corrected two-tailed Wilcoxon signed-rank tests; significance level equates to α /n (α =0.05), in which n is the number of tests. Significance is indicated in all figures below the respective boxplot by: (ns) not significant; (*) p< 0.05/n. For comparison between two groups, which did not violate the assumptions of normality (Shapiro-Wilk test) and homogeneity of variance (Bartlett’s test) were analyzed with two-tailed unpaired t-test, otherwise two-tailed Mann-Whitney test. Significance is indicated in all figures above boxplots by: (ns) not significant; (*) p<0.05. For statistical tests involving two factors, two-way ANOVAs were applied, followed by planned, pairwise multiple comparisons (Bonferroni *post-hoc* pairwise comparisons); significance is indicated in all figures above boxplots by: lowercase letters indicate differences between groups (p<0.05) or (ns) not significant. Respective statistical tests used, sample sizes, and descriptive statistics can be found in Supplemental Table S1, S2 and S3 for main figures and Supporting Information figures. Data were presented as Tukey box plots, with 50% of the values being located within the boxes and whiskers representing 1.5 interquartile range. Outsiders were indicated as open circles. The median was indicated as a bold line and the mean as a cross within the box plot. Unless stated otherwise, experiments had a sample size of 16. Figure alignments were performed with *Adobe Photoshop CC* 2019 and *Adobe Illustrator CC* 2019.

