## Supplementary material for "Insulin signaling gates long-term memory formation in *Drosophila* larvae": S1 Table

### S1 Table: Sample size, mean±s.e.m. and statistical details of test against chance level.

|  | Group^1^ | n | Mean±s.e.m | Normally  distributed^2^ | Shapiro-Wilk test | p-value^3^ | 𝛼^4^ | Significant^5^ |
| --- | --- | --- | --- | --- | --- | --- | --- | --- |
| Fig 1C | 1 | 15 | -0.198±0.030 | yes | W=0.894, p=0.076 | <0.0001 | 0.0125 | * |
|  | 2 | 13 | -0.228±0.055 | yes | W=0.953, p=0.642 | 0.001 |  | * |
|  | 3 | 14 | -0.223±0.023 | yes | W=0.965, p=0,798 | <0.0001 |  | * |
|  | 4 | 12 | -0.006±0.044 | yes | W=0.920, p=0.282 | 0.898 |  | ns |
| Fig 2A | 1 | 16 | -0.238±0.040 | no | W=0.853, p=0.015 | 0.0005 | 0.0125 | * |
|  | 2 | 16 | -0.002±0.035 | yes | W=0.939, p=0.335 | 0.838 |  | ns |
|  | 3 | 16 | -0.245±0.043 | yes | W=0.943, p=0.381 | 0.0002 |  | * |
|  | 4 | 18 | -0.166±0.048 | no | W=0.865, p=0.015 | 0.003 |  | * |
| Fig 2B | 1 | 14 | -0.218±0.036 | yes | W=0.982, p=0.986 | <0.0001 | 0.0125 | * |
|  | 2 | 14 | -0.038±0.033 | yes | W=0.909, p=0.153 | 0.275 |  | ns |
|  | 3 | 14 | -0.067±0.047 | yes | W=0.977, p=0.953 | 0.176 |  | ns |
|  | 3 | 14 | -0.028±0.026 | yes | W=0.981, p=0.978 | 0.312 |  | ns |
| Fig 2C | 1 | 11 | -0.309±0.040 | yes | W=0.950, p=0.646 | <0.0001 | 0.0125 | * |
|  | 2 | 12 | -0.271±0.023 | yes | W=0.977, p=0.966 | <0.0001 |  | * |
|  | 3 | 15 | -0.347±0.028 | yes | W=0.925, p=0.226 | <0.0001 |  | * |
|  | 3 | 16 | -0.079±0.035 | yes | W=0.944, p=0.40 | 0.041 |  | ns |
| Fig 3A | 1 | 15 | -0.187±0.035 | yes | W=0.984, p=0.991 | 0.0001 | 0.008 | * |
|  | 2 | 14 | -0.201±0.033 | yes | W=0.948, p=0.523 | <0.0001 |  | * |
|  | 3 | 14 | -0.174±0.033 | yes | W=0.977, p=0.952 | 0.0002 |  | * |
|  | 4 | 14 | -0.054±0.035 | yes | W=0.951, p=0.580 | 0.149 |  | ns |
|  | 5 | 14 | -0.038±0.050 | yes | W=0.887, p=0.102 | 0.469 |  | ns |
|  | 6 | 18 | -0.218±0.040 | yes | W=0.982, p=0.964 | <0.0001 |  | * |
| Fig 4B | @ 30 min |  |  |  |  |  |  |  |
|  | 1 | 16 | -0.333±0.035 | yes | W=0.967, p=0.789 | <0.0001 | 0.0125 | * |
|  | 2 | 8 | -0.266±0.021 | yes | W=0.921, p=0.441 | <0.0001 |  | * |
|  | 3 | 16 | -0.354±0.026 | yes | W=0.921, p=0.175 | <0.0001 |  | * |
|  | 4 | 12 | -0.218±0.030 | yes | W=0.965, p=0.850 | <0.0001 |  | * |
|  | @ 60 min |  |  |  |  |  |  |  |
|  | 1 | 16 | -0.290±0.030 | yes | W=0.930, p=0.244 | <0.0001 | 0.0125 | * |
|  | 2 | 14 | -0.286±0.025 | yes | W=0.974, p=0.929 | <0.0001 |  | * |
|  | 3 | 16 | -0.131±0.033 | yes | W=0.949, p=0.467 | <0.0001 |  | * |
|  | 4 | 14 | -0.109±0.037 | yes | W=0.910, p=0.160 | 0.011 |  | * |
| S1A Fig | @ 15 min |  |  |  |  |  |  |  |
|  | 1 | 16 | 0.000±0.154^5^ | yes | W=0.906, p=0.10 | >0.999 | 0.025 | ns |
|  | 2 | 16 | 0.833±0.140 | yes | W=0.958, p=0.630 | <0.0001 |  | * |
|  | @ 30 min |  |  |  |  |  |  |  |
|  | 1 | 16 | 0.000±0.194^5^ | yes | W=0.929, p=0.233 | 0.980 | 0.025 | ns |
|  | 2 | 16 | 1.623±0.271 | no | W=0.815, p=0.004 | <0.0001 |  | * |
|  | @ 60 min |  |  |  |  |  |  |  |
|  | 1 | 16 | 0.000±0.097^5^ | yes | W=0.985, p=0.989 | 0.940 | 0.025 | ns |
|  | 2 | 16 | 0.161±0.109 | no | W=0.885, p=0.046 | 0.376 |  | ns |
| S1C Fig | AM |  |  |  |  |  |  |  |
|  | 1 | 16 | 0.379±0.043 | yes | W=0.950, p=0.491 | <0.0001 | 0.025 | * |
|  | 2 | 16 | 0.289±0.050 | yes | W=0.989, p=0.999 | <0.0001 |  | * |
|  | BA |  |  |  |  |  |  |  |
|  | 1 | 16 | 0.284±0.063 | yes | W=0.966, p=0.761 | 0.0009 | 0.025 | * |
|  | 2 | 16 | 0.283±0.055 | no | W=0.877, p=0.035 | 0.0017 |  | * |
|  | salt |  |  |  |  |  |  |  |
|  | 1 | 16 | -0.393±0.050 | yes | W=0.962, p=0.697 | <0.0001 | 0.025 | * |
|  | 2 | 16 | -0.524±0.064 | yes | W=0.942, p=0.373 | <0.0001 |  | * |
| S2A Fig | 1 | 14 | -0.190±0.038 | yes | W=0.974, p=0.926 | 0.0002 | 0.0125 | * |
| continuation | | | | | | | | |
|  | **Group^1^** | **n** | **Mean±s.e.m** | **Normally**  **distributed^2^** | **Shapiro-Wilk test** | **p-value^3^** | **𝛼^4^** | **Significant^5^** |
| S2A Fig | 2 | 15 | -0.165±0.044 | yes | W=0.952, p=0.553 | 0.002 |  | * |
|  | 3 | 14 | -0.154±0.038 | yes | W=0.946, p=0.499 | 0.001 |  | * |
|  | 4 | 15 | -0.178±0.045 | yes | W=0.973, p=0.896 | 0.002 |  | * |
| S2B Fig | 1 | 10 | -0.271±0.049 | yes | W=0.957, p=0.712 | 0.0004 | 0.0125 | * |
|  | 2 | 8 | -0.246±0.070 | yes | W=0.930, p=0.516 | 0.009 |  | * |
|  | 3 | 9 | -0.322±0.046 | yes | W=0.937, p=0.552 | 0.0001 |  | * |
|  | 4 | 9 | -0.253±0.051 | yes | W=0.939, p=0.567 | 0.001 |  | * |
| S3B Fig | 1 | 16 | 0.330±0.063 | yes | W=0.970, p=0.833 | 0.0001 | 0.008 | * |
|  | 2 | 16 | 0.403±0.075 | yes | W=0.914, p=0.133 | <0.0001 |  | * |
|  | 3 | 16 | 0.491±0.062 | yes | W=0.931, p=0.255 | <0.0001 |  | * |
|  | 4 | 16 | 0.399±0.061 | no | W=0.809, p=0.004 | <0.0001 |  | * |
|  | 5 | 16 | 0.407±0.047 | yes | W=0.954, p=0.558 | <0.0001 |  | * |
|  | 6 | 16 | 0.442±0.043 | yes | W=0.949, p=0.478 | <0.0001 |  | * |
| S3C Fig | 1 | 16 | 0.180±0.043 | yes | W=0.915, p=0.141 | 0.0002 | 0.008 | * |
|  | 2 | 16 | 0.085±0.036 | no | W=0.860, p=0.019 | 0.016 |  | ns |
|  | 3 | 16 | 0.116±0.029 | yes | W=0.965, p=0.754 | 0.002 |  | * |
|  | 4 | 16 | 0.179±0.046 | yes | W=0.924, p=0.197 | 0.0006 |  | * |
|  | 5 | 16 | 0.134±0.039 | no | W=0.806, p=0.003 | 0.0005 |  | * |
|  | 6 | 15 | 0.135±0.043 | yes | W=0.964, p=0.765 | 0.007 |  | * |
| S3D Fig | 1 | 23 | -0.383±0.063 | yes | W=0.981, p=0.919 | <0.0001 | 0.008 | * |
|  | 2 | 23 | -0.372±0.050 | yes | W=0.943, p=0.211 | <0.0001 |  | * |
|  | 3 | 22 | -0.361±0.064 | yes | W=0.943, p=0.237 | <0.0001 |  | * |
|  | 4 | 23 | -0.323±0.043 | yes | W=0.933, p=0.128 | <0.0001 |  | * |
|  | 5 | 22 | -0.419±0.055 | yes | W=0.954, p=0.378 | <0.0001 |  | * |
|  | 6 | 22 | -0.350±0.056 | yes | W=0.984, p=0.963 | <0.0001 |  | * |
| S3F Fig | 1 | 8 | -0.058±0.118 | yes | W=0.974, p=0.923 | 0.637 | 0.025 | ns |
|  | 2 | 8 | -0.087±0.231 | yes | W=0.950, p=0.729 | 0.719 |  | ns |
| S3G Fig | 1 | 8 | 0.000±0.244 | yes | W=0.866, p=0.137 | >0.999 | 0.025 | ns |
|  | 2 | 8 | -0.243±0.114 | yes | W=0.890, p=0.235 | 0.069 |  | ns |
| S3H Fig | 1 | 7 | 0.000±0.142 | yes | W=0.942, p=0.657 | >0.999 | 0.025 | ns |
|  | 2 | 7 | -0.327±0.128 | yes | W=0.952, p=0.744 | 0.044 |  | * |
| S4A Fig | 1 | 10 | -0.181±0.060 | yes | W=0.989, p=0.994 | 0.016 | 0.025 | * |
|  | 2 | 9 | -0.101±0.039 | yes | W=0.930, p=0.443 | 0.020 |  | * |
| S4B Fig | 1 | 12 | -0.207±0.045 | yes | W=0.950, p=0.634 | 0.0007 | 0.025 | * |
|  | 2 | 12 | -0.253±0.035 | yes | W=0.932, p=0.402 | <0.0001 |  | * |
| S4C Fig | 1 | 16 | 0.000±0.146 | yes | W=0.893, p=0.063 | >0.999 | 0.025 | ns |
|  | 2 | 16 | 0.173±0.116 | yes | W=0.964, p=0.725 | 0.155 |  | ns |

^1^Numbers correspond to boxplots from left to right. ^2^Significance level was set to 𝛼=0.05, p≥0.05 indicates normal distribution. One-sample t-test or Wilcoxon signed-rank test. Theoretical mean was 0.000. ^3^Bonferroni-corrected significance level equates to 𝛼/n (𝛼=0.05). ^4^ns indicates p≥0.05/n, * indicates p<0.05/n. ^5^Normalized to group 1.
