## Supplementary material for "Insulin signaling gates long-term memory formation in *Drosophila* larvae": S2 Table

### S2 Table: Statistical details of unpaired t-test or Mann-Whitney test.

|  | Statistical test | Statistical description | p-value | Significant^2^ |
| --- | --- | --- | --- | --- |
| S1C Fig | AM |  |  |  |
|  | Unpaired t-test^1^ | t=1.365, df=30 | 0.183 | ns |
|  | BA |  |  |  |
|  | Mann-Whitney test^1^ | U=122.5 | 0.846 | ns |
|  | Salt |  |  |  |
|  | Unpaired t-test^1^ | t=1.617, df=30 | 0.116 | ns |
| S4A Fig | Unpaired t-test^1^ | t=1.031, df=17 | 0.317 | ns |
| S4B Fig | Unpaired t-test^1^ | t=0.801, df=22 | 0.432 | ns |

^1^Two-tailed. ^2^Significance level was set to 𝛼=0.05. ns indicates p≥0.05, * indicates p<0.05.
