## Supplementary material for "Insulin signaling gates long-term memory formation in *Drosophila* larvae": S3 Table

### S3 Table: Statistical details of Two-way ANOVA.

|  | Two-way ANOVA  (Factors) | Statistical description of main effects^1^ | Significant^2^ | Bonferroni *post-hoc* pairwise comparisons^3^ |
| --- | --- | --- | --- | --- |
| Fig 1C | Cold shock | F(1,50)=6.556, p=0.014 | * | 1 vs 2, p>0.9999, ns |
|  | Sucrose consumption | F(1,50)=7.220, p=0.010 | * | 1 vs 3, p>0.9999, ns |
|  | Interaction | F(1,50)=10.95, p=0.002 | * | 1 vs 4, p=0.003, * |
|  |  |  |  | 2 vs 3, p>0.9999, ns |
|  |  |  |  | 2 vs 4, p=0.001, * |
|  |  |  |  | 3 vs 4, p=0.0009, ns |
| Fig 2A | Genotype | F(1,62)=14.09, p=0.0004 | * | 1 vs 2, 0.0012, * |
|  | Sucrose consumption | F(1,62)=4.234, p=0.044 | * | 1 vs 3, >0.9999, ns |
|  | Interaction | F(1,62)=3.62, p=0.062 | ns | 1 vs 4, >0.9999, ns |
|  |  |  |  | 2 vs 3, 0.0009, * |
|  |  |  |  | 2 vs 4, 0.036, * |
|  |  |  |  | 3 vs 4, p>0.9999, ns |
| Fig 2B | Genotype | F(1,52)=9.151, p=0.004 | * | 1 vs 2, p=0.006, * |
|  | Sucrose consumption | F(1,52)=4.954, p=0.030 | * | 1 vs 3, p=0.029, * |
|  | Interaction | F(1,52)=3.765, p=0.058 | ns | 1 vs 4, p=0.003, * |
|  |  |  |  | 2 vs 3, p=0.994, ns |
|  |  |  |  | 2 vs 4, p>0.9999, ns |
|  |  |  |  | 3 vs 4, 0.9712, ns |
| Fig 2C | Genotype | F(1,50)=21.81, p<0.0001 | * | 1 vs 2, p>0.9999, ns |
|  | Sucrose consumption | F(1,50)=5.490, p=0.023 | * | 1 vs 3, p>0.9999, ns |
|  | Interaction | F(1,50)=12.37, p=0.0009 | * | 1 vs 4, p<0.0001, * |
|  |  |  |  | 2 vs 3, p=0.626, ns |
|  |  |  |  | 2 vs 4, p=0.0006, * |
|  |  |  |  | 3 vs 4, p<0.0001, * |
| Fig 3A | Genotype | F(2,83)=4.555, p=0.013 | * | 1 vs 2, p>0.9999, ns |
|  | Cold shock | F(2,83)=11.88, p=0.0009 | * | 1 vs 3, p>0.9999, ns |
|  | Interaction | F(2,83)=6.916, p=0.002 | * | 1 vs 4, p=0.0.276, ns |
|  |  |  |  | 1 vs 5, p=0.002, * |
|  |  |  |  | 1 vs 6, p>0.9999, ns |
|  |  |  |  | 2 vs 3, p>0.9999, ns |
|  |  |  |  | 2 vs 4, p=0.002, * |
|  |  |  |  | 2 vs 5, p=0.0008, * |
|  |  |  |  | 2 vs 6, p>0.9999, ns |
|  |  |  |  | 3 vs 4, p=0.540, ns |
|  |  |  |  | 3 vs 5, p=0.005, * |
|  |  |  |  | 3 vs 6, p>0.9999, ns |
|  |  |  |  | 4 vs 5, p>0.9999, ns |
|  |  |  |  | 5 vs 6, p<0.0001, * |
| Fig 4B | @ 30 min |  |  |  |
|  | Cycloheximide treatment | F(1,48)=9.959, p=0.003 | * | 1 vs 2, p>0.9999, ns |
|  | Sucrose consumption | F(1,48)=0.178, p=0.675 | ns | 1 vs 3, p>0.9999, ns |
|  | Interaction | F(1,48)=1.13, p=0.293 | ns | 1 vs 4, p=0.057, ns |
|  |  |  |  | 2 vs 3, p=0.447, ns |
|  |  |  |  | 2 vs 4, p>0.9999, ns |
|  |  |  |  | 3 vs 4, 0.015, * |
|  | @ 60 min |  |  |  |
|  | Cycloheximide treatment | F(1,56)=9.981, p=0.003 | * | 1 vs 2, p>0.9999, ns |
|  | Sucrose consumption | F(1,56)=6.854, p=0.011 | * | 1 vs 3, p>0.9999, ns |
|  | Interaction | F(1,56)=9.244, p=0.004 | * | 1 vs 4, p=0.0003, * |
|  |  |  |  | 2 vs 3, p>0.9999, ns |
|  |  |  |  | 2 vs 4, p=0.0003, * |
|  |  |  |  | 3 vs 4, p=0.002, * |
| S2A Fig | Cold shock | F(1,54)=0.0004, p=0.985 | ns | 1 vs 2, p=0.931, ns |
|  | Sucrose consumption | F(1,54)=0.0752, p=0.785 | ns | 1 vs 3, p=0.973, ns |
|  | Interaction | F(1,54)=0.348, p=0.558 | ns | 1 vs 4, p=0.997, ns |
|  |  |  |  | 2 vs 3, p=0.998, ns |
| continuation | | | | |
|  | Two-way ANOVA  (Factors) | Statistical description of main effects^1^ | Significant^2^ | Bonferroni *post-hoc* pairwise comparisons^3^ |
| S2A Fig |  |  |  | 2 vs 4, p=0.997, ns |
|  |  |  |  | 3 vs 4, p=0.996, ns |
| S2B Fig | Cold shock | F(1,32)=0.7712, p=0.386 | ns | 1 vs 2, p=0.899, ns |
|  | Hypercaloric food | F(1,32)=0.2887, p=0.595 | ns | 1 vs 3, p=0.988, ns |
|  | Interaction | F(1,32)=0.1729, p=0.680 | ns | 1 vs 4, p=0.995, ns |
|  |  |  |  | 2 vs 3, p=0.765, ns |
|  |  |  |  | 2 vs 4, p=0.796, ns |
|  |  |  |  | 3 vs 4, p>0.9999, ns |
| S3B Fig | Genotype | F(2,90)=1.521, p=0.224 | ns | 1 vs 2, p>0.9999, ns |
|  | Sucrose | F(1,90)=0.0330, p=0.856 | ns | 1 vs 3, p>0.9999, ns |
|  | Interaction | F(2,90)=0.4962, p=0.612 | ns | 1 vs 4, p>0.9999, ns |
|  |  |  |  | 1 vs 5, p=0.857, ns |
|  |  |  |  | 1 vs 6, p>0.9999, ns |
|  |  |  |  | 2 vs 3, p>0.9999, ns |
|  |  |  |  | 2 vs 4, p>0.9999, ns |
|  |  |  |  | 2 vs 5, p>0.9999, ns |
|  |  |  |  | 2 vs 6, p>0.9999, ns |
|  |  |  |  | 3 vs 4, p>0.9999, ns |
|  |  |  |  | 3 vs 5, p>0.9999, ns |
|  |  |  |  | 3 vs 6, p>0.9999, ns |
|  |  |  |  | 4 vs 5, p>0.9999, ns |
|  |  |  |  | 4 vs 6, p>0.9999, ns |
|  |  |  |  | 5 vs 6, p>0.9999, ns |
| S3C Fig | Genotype | F(2,89)=1.736, p=0.182 | ns | 1 vs 2, p>0.9999, ns |
|  | Sucrose | F(1,89)=0.4836, p=0.489 | ns | 1 vs 3, p>0.9999, ns |
|  | Interaction | F(2,89)=0.2097, p=0.811 | ns | 1 vs 4, p>0.9999, ns |
|  |  |  |  | 1 vs 5, p>0.9999, ns |
|  |  |  |  | 1 vs 6, p>0.9999, ns |
|  |  |  |  | 2 vs 3, p>0.9999, ns |
|  |  |  |  | 2 vs 4, p>0.9999, ns |
|  |  |  |  | 2 vs 5, p>0.9999, ns |
|  |  |  |  | 2 vs 6, p>0.9999, ns |
|  |  |  |  | 3 vs 4, p>0.9999, ns |
|  |  |  |  | 3 vs 5, p>0.9999, ns |
|  |  |  |  | 3 vs 6, p>0.9999, ns |
|  |  |  |  | 4 vs 5, p>0.9999, ns |
|  |  |  |  | 4 vs 6, p>0.9999, ns |
| continuation | | | | |
|  | **Two-way ANOVA**  **(Factors)** | **Statistical description of main effects^1^** | **Significant^2^** | **Bonferroni *post-hoc* pairwise comparisons^3^** |
| S3C Fig |  |  |  | 5 vs 6, p>0.9999, ns |
| S3D Fig | Genotype | F(2,129)=0.3663, p=0.694 | ns | 1 vs 2, p>0.9999, ns |
|  | Sucrose | F(1,129)=0.0359, p=0.850 | ns | 1 vs 3, p>0.9999, ns |
|  | Interaction | F(2,129)=0.467, p=0.628 | ns | 1 vs 4, p>0.9999, ns |
|  |  |  |  | 1 vs 5, p>0.9999, ns |
|  |  |  |  | 1 vs 6, p>0.9999, ns |
|  |  |  |  | 2 vs 3, p>0.9999, ns |
|  |  |  |  | 2 vs 4, p>0.9999, ns |
|  |  |  |  | 2 vs 5, p>0.9999, ns |
|  |  |  |  | 2 vs 6, p>0.9999, ns |
|  |  |  |  | 3 vs 4, p>0.9999, ns |
|  |  |  |  | 3 vs 5, p>0.9999, ns |
|  |  |  |  | 3 vs 6, p>0.9999, ns |
|  |  |  |  | 4 vs 5, p>0.9999, ns |
|  |  |  |  | 4 vs 6, p>0.9999, ns |
|  |  |  |  | 5 vs 6, p>0.9999, ns |

^1^F(DFn, Dfd), p-value. ^2^Significance level was set to 𝛼=0.05. ns indicates p≥0.05, * indicates p<0.05. ^3^Numbers correspond to boxplots from left to right. Adjusted p-values after Bonferroni *post-hoc* pairwise comparison. Significance level was set to 𝛼=0.05, ns indicates p≥0.05, * indicates p<0.05.
