## Supplementary material for "Insulin signaling gates long-term memory formation in *Drosophila* larvae": S1 Fig

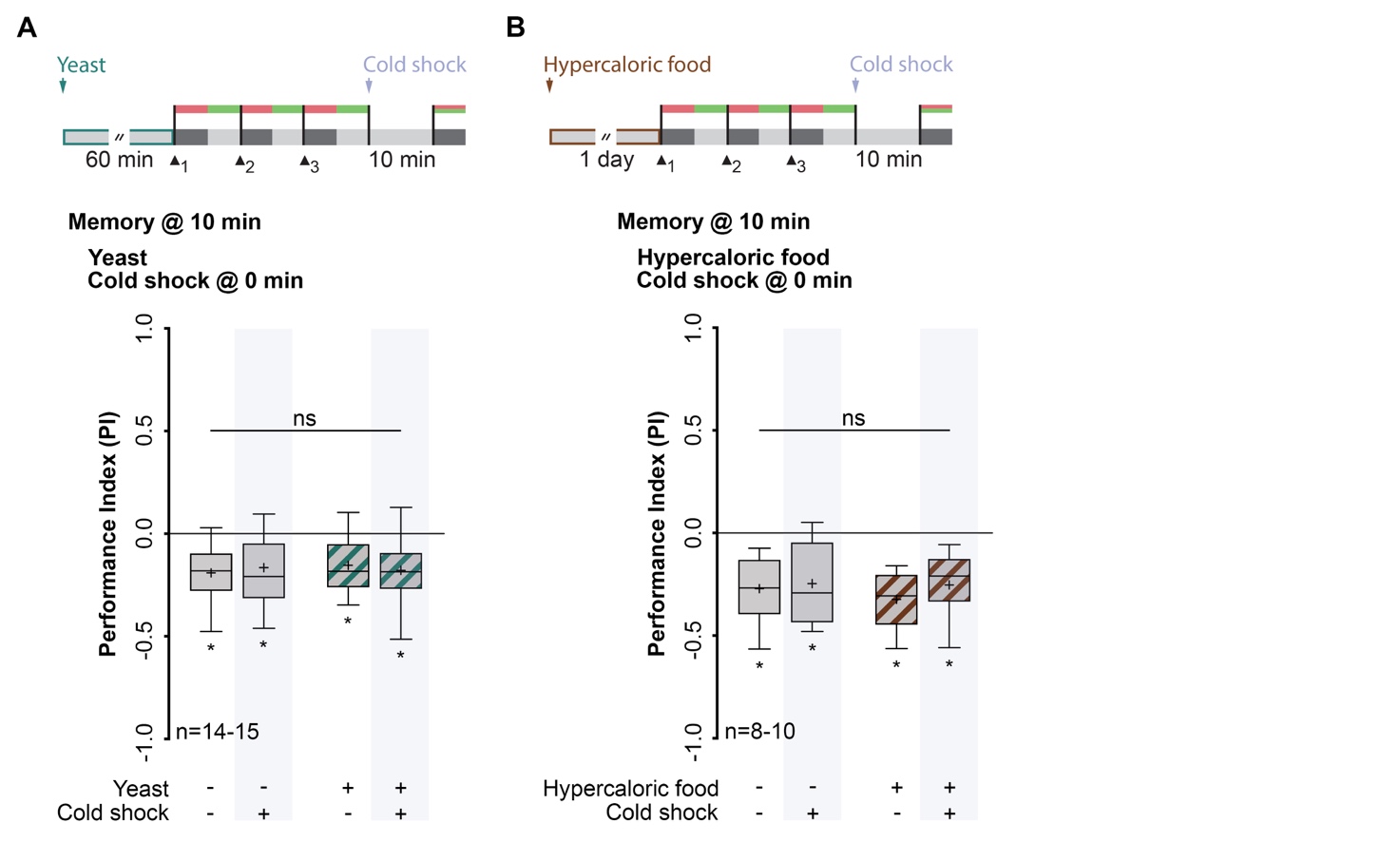

### S1 Fig. Time-dependent sucrose feeding in wild-type larvae and task-relevant sensory-motor abilities.

(A) Top: Sucrose consumption quantification using a photometer-quantified dye feeding assay. Wild-type larvae fed either on dye alone (left) or sucrose + dye (right) for 15, 30, or 60 min. By normalizing to the dye-only group a Relative Consumption of Sucrose Index (R.C.) was calculated. Bottom: Sucrose consumption is dependent on the satiation state of larvae. An increased sucrose consumption was observed at 15 and 30 min, whereas sucrose consumption at 60 min was similar to dye-only group. (B) Representative image showing wild-type larvae fed for 15 min, 30 or 60 min either on dye or on sucrose + dye (from left, in order). (C) Top: Odor preference and high-salt avoidance assays after ingesting sucrose for 60 min. Naïve AM preference left, naïve BA preference middle, salt avoidance right. Olfactory perception was analyzed by calculating an Olfactory Preference Index (PREF). High salt avoidance was analyzed by calculating a Gustatory Avoidance Index (GAI). Bottom: Task-relevant sensory-motor abilities were not altered after sucrose consumption. Sucrose consumption, naïve odor preference, and high salt avoidance above the level of chance was tested using Bonferroni-corrected one-sample t-tests or Wilcoxon signed-rank test (ns p≥0.025; * p<0.025; 𝛼=0.025). Differences between groups in (B) were determined using unpaired t-test or Mann-Whitney test. Statistically non-significant differences between groups (p≥0.05) are indicated as ns. For more statistical details see also S1 Table and S2 Table. Data are shown as Tukey box plots; line, median; cross, mean; box, 75th-25th percentiles; whiskers, 1.5 interquartile range; small circles, outlier (n≥8). AM, n-amyl acetate; BA, benzaldehyde.
