## Supplementary material for "Insulin signaling gates long-term memory formation in *Drosophila* larvae": S2 Fig

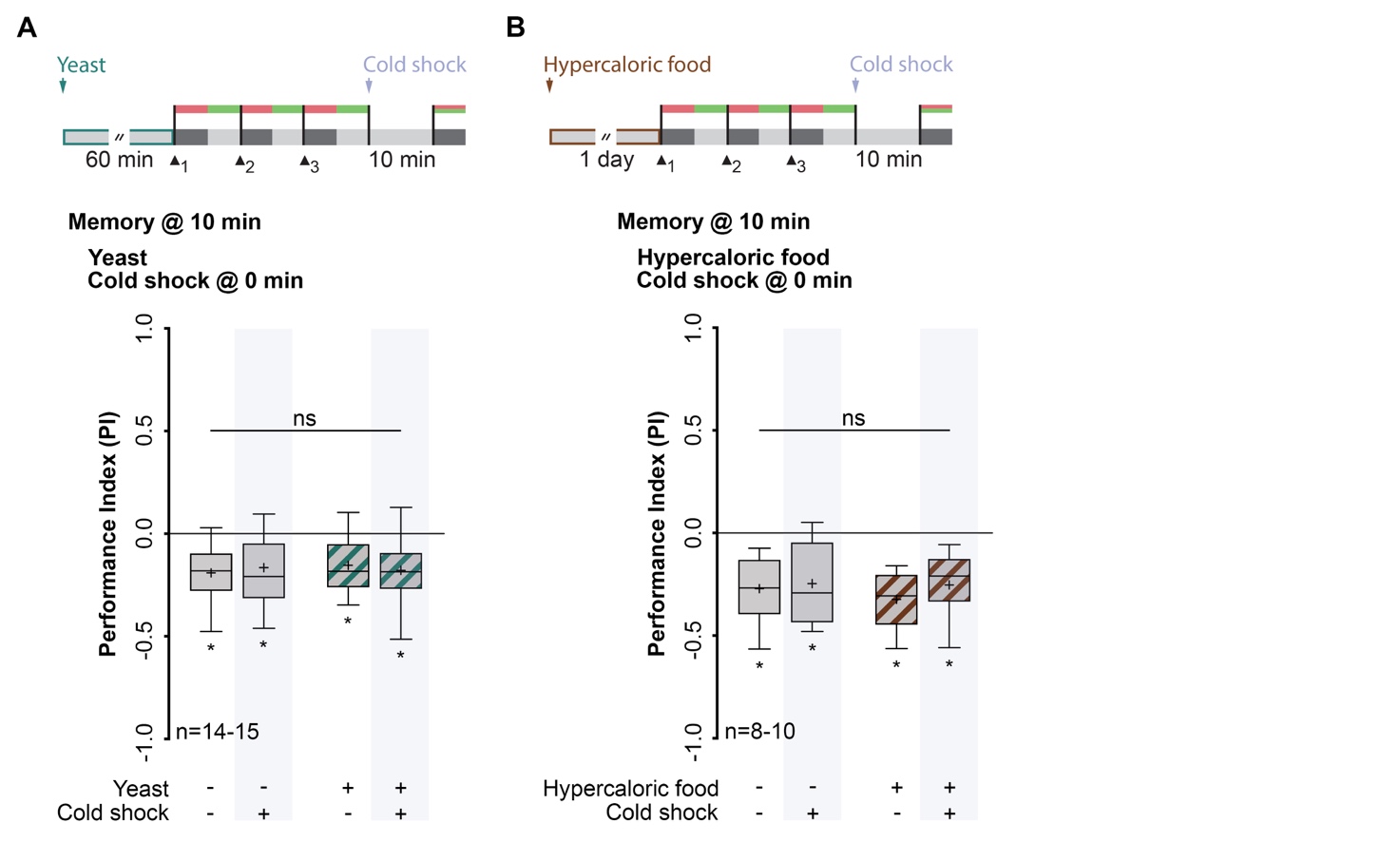

### S2 Fig. Aversive olfactory memory formation after a protein-rich diet.

(A) Top: Training and treatment protocols. Larvae fed for 60 min on yeast and identification of lARM was carried out by applying a cold shock directly after training. Memory was tested 10 min after training. Bottom: Feeding on yeast does not inhibit lARM formation in wild-type larvae. (B) Top: Training and treatment protocols. Larvae were kept for one day on protein and carbohydrate rich food (hypercaloric food). Identification of lARM was carried out by applying a cold shock directly after training. Memory was tested 10 minutes after training. Bottom: Feeding for one day on hypercaloric food does not inhibit lARM formation in wild-type larvae. Memory performance above the level of chance was tested using Bonferroni-corrected one-sample t-tests (ns p≥0.0125; * p<0.0125; 𝛼=0.0125). Differences between groups were determined using two-way ANOVA followed by Bonferroni *post-hoc* pairwise comparisons. Statistically non-significant differences between groups (p≥0.05) are indicated as ns. For more statistical details see also Table S1Table and S3 Table. Data are shown as Tukey box plots; line, median; cross, mean; box, 75th-25th percentiles; whiskers, 1.5 interquartile range; small circles, outlier (n≥8). lARM, larval anesthesia resistant memory.
