## Supplementary material for "Insulin signaling gates long-term memory formation in *Drosophila* larvae": S3 Fig

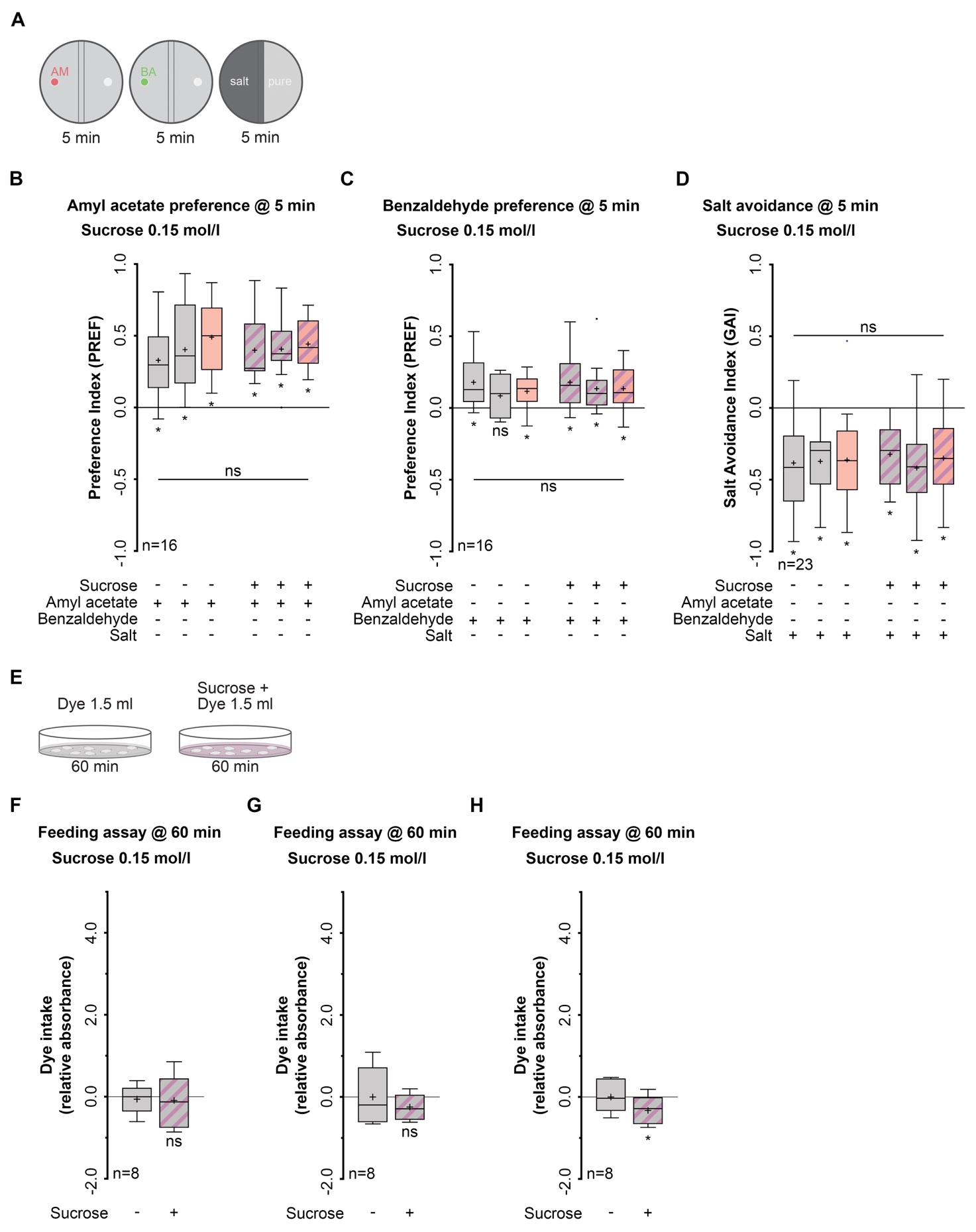

### S3 Fig. Task-relevant sensory-motor abilities and sucrose feeding in larvae expressing the dominant negative form of the insulin receptor (InR^DN^).

(A) Odor preference and high-salt avoidance assays after ingesting sucrose for 60 min. Naïve amyl acetate (AM) left preference left, naïve benzaldehyde (BA) preference middle, salt avoidance right. Olfactory perception was analyzed by calculating an Olfactory Preference Index (PREF). High salt avoidance was analyzed by calculating a Gustatory Avoidance Index (GAI). (B)-(D) Task-relevant sensory-motor abilities are not altered in larvae expressing a dominant negative form of the insulin receptor (InR^DN^) in the MB KCs via the OK107 driver line. (B) naïve odor preference for AM. (C) naïve odor preference for BA. (D) naïve avoidance of a high salt concentration. Naïve odor preference and high salt avoidance above the level of chance was tested using Bonferroni-corrected one-sample t-tests (ns p≥0.008; * p<0.008; 𝛼=0.008). Differences between the groups were determined using two-way ANOVA followed by Bonferroni *post-hoc* pairwise comparisons. Statistically non-significant differences between groups (p≥0.05) are indicated as ns. For more statistical details see also Table S1Table and S3 Table. (E) Sucrose consumption quantification using a photometer-quantified dye feeding assay. A Relative Consumption of Sucrose Index (R.C.) was calculated by normalizing to the dye-only control. (F)-(H) Expression of a dominant negative insulin receptor (*InR^DN^*) transgene in MB KCs via OK107 driver line leads to a small reduction in sucrose consumption. F: Sucrose consumption in OK107/+ control group. G: Sucrose consumption in UAS-*InR^DN^*/+ control group. G: Sucrose consumption in OK107/UAS-*InR^DN^* experimental group. Sucrose consumption above the level of chance was tested using Bonferroni-corrected one-sample t-tests. For (ns p≥0.025; * p<0.025; 𝛼=0.025). For more statistical details see also Table S1Table. Data are shown as Tukey box plots; line, median; cross, mean; box, 75th-25th percentiles; whiskers, 1.5 interquartile range; small circles, outlier (n≥8). DN, dominant negative; InR, insulin receptor; KC, Kenyon cell; lARM, larval anesthesia resistant memory; UAS, upstream activation sequence.
