## Supplementary material for "Insulin signaling gates long-term memory formation in *Drosophila* larvae": S4 Fig

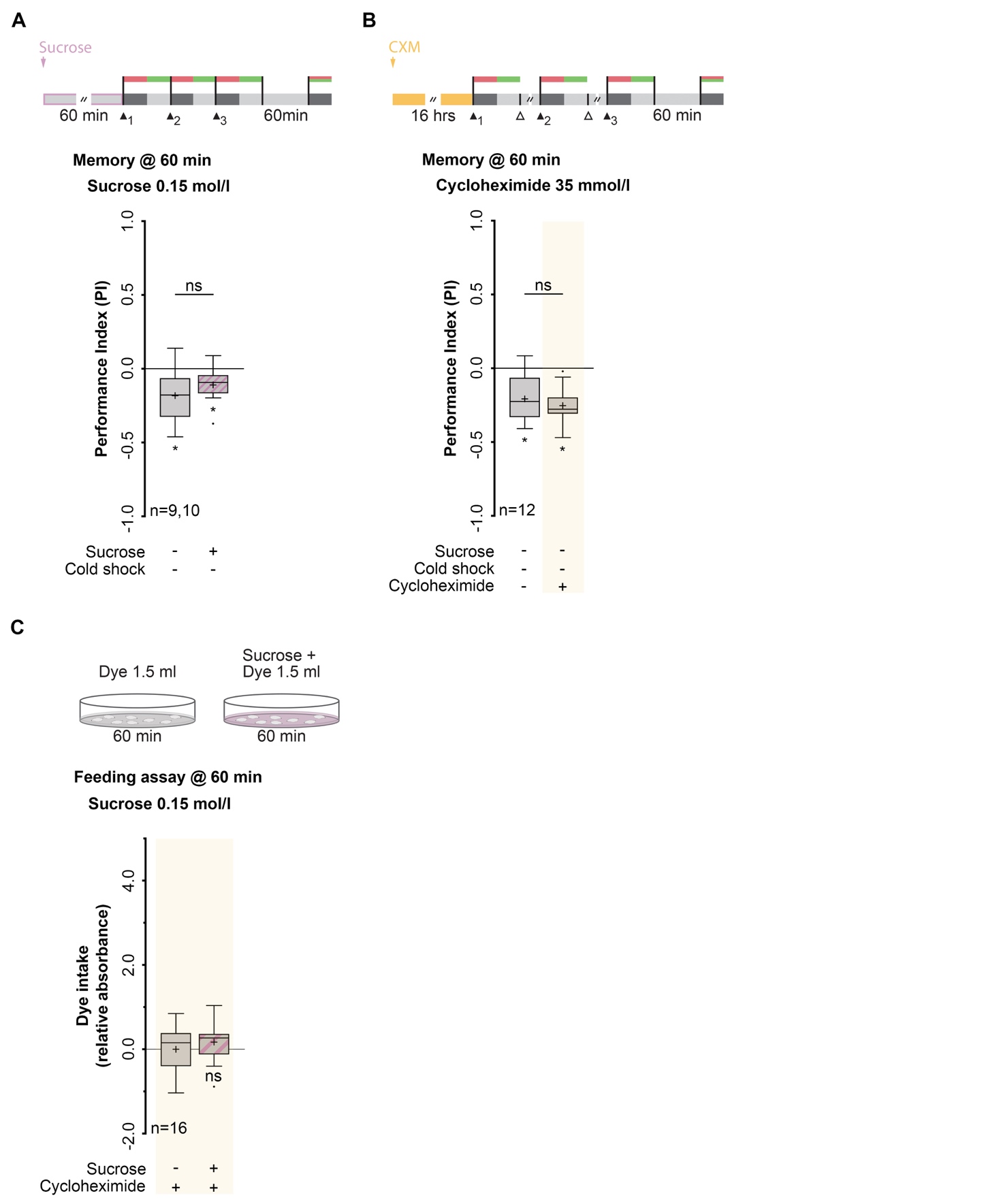

### S4 Fig. Stability of aversive olfactory memory after sucrose consumption and formation of aversive olfactory memory after spaced training.

(A) Top: Training and different treatment protocols. Memory was tested 60 min after training. Bottom: Memory after sucrose consumption was statistically indistinguishable from lARM formed after being fed on tap water. (B) Top: Training and different treatment protocols. Wild-type larvae were fed CXM for 16 hours. Odor-high salt conditioning was performed using a spaced training protocol consisting of three cycles separated by 15 min rest intervals (open triangle). Memory was tested 60 min after training. Bottom: The application of CXM did not affect the aversive olfactory memory that was formed after a spaced training protocol. (C) Top: Sucrose consumption quantification in wild-type larvae using a photometer-quantified dye feeding assay. A Relative Consumption of Sucrose Index (R.C.) was calculated by normalizing to the dye-only control. Both groups were fed CXM for 16 hours before feeding on sucrose. Bottom: Sucrose consumption is not altered after CXM treatment. Memory performance and sucrose consumption above the level of chance was tested using Bonferroni-corrected one-sample t-tests. For (ns p≥0.025; * p<0.025; 𝛼=0.025). Differences between groups in (A) and (B) were determined using unpaired t-test or Mann-Whitney test. Statistically non-significant differences between groups (p≥0.05). For more statistical details see also Table S1 and Table S2. Data are shown as Tukey box plots; line, median; cross, mean; box, 75th-25th percentiles; whiskers, 1.5 interquartile range; small circles, outlier (n≥8). CXM, cycloheximide; lARM, larval anesthesia resistant memory.
